## Supplementary materials for "DWV 3C protease uncovers the diverse catalytic triad in insect RNA viruses"

**Supplementary Table 1 List of primers used in this study**

| | |
|---|---|
| GST-3C$^{pro}$-5-BamHI | TTTGGATCCGGATTGAAATATAGTGAAGCAGT |
| GST-3C$^{pro}$-3-NotI | TTTGCGGCCGCTTCACTCTCGATTGCTTTACC |
| pGEX 5' primer | GGGCTGGCAAGCCACGTTTGGTG |
| pGEX 3' primer | CCGGGAGCTGCATGTGTCAGAGG |
| N2134A-For | AATAAAATTTTACAGGCGATGGTTTACATTGGT |
| N2134A-Rev | ACCAATGTAAACCATCGCCTGTAAAATTTTATT |
| R2156A-For | CGAGATATTAATTTTGCGTGTCTTATGCTTCAT |
| R2156A-Rev | ATGAAGCATAAGACACGCAAAATTAATATCTCG |
| L2168A-For | AGGCAATGTTTAATGGCGAGGCATTATCTCGAG |
| L2168A-Rev | CTCGAGATAATGCCTCGCCATTAAACATTGCCT |
| H2170A-For | TGTTTAATGTTAAGGGCGTATCTCGAGTCAACT |
| H2170A-Rev | AGTTGACTCGAGATACGCCCTTAACATTAAACA |
| Y2171A-For | TTAATGTTAAGGCATGCGCTCGAGTCAACTGCC |
| Y2171A-Rev | GGCAGTTGACTCGAGCGCATGCCTTAACATTAA |
| H2190A-For | TATTTTAAGTATATTGCGAATCAAGAGACTAGA |
| H2190A-Rev | TCTAGTCTCTTGATTCGCAATATACTTAAAATA |
| D2225A-For | GGAGAGGAGTCGTTTGCGAGTAATATCGTGCTT |
| D2225A-Rev | AAGCACGATATTACTCGCAAACGACTCCTCTCC |
| N2227A-For | GAGTCATTTGATAGCGCGATCGTGCTTGTGACT |
| N2227A-Rev | AGTCACAAGCACGATCGCGCTATCAAATGACTC |
| I2283A-For | AAAACTCCAATAAGCGCGAACGCTGATGGTTTG |
| I2283A-Rev | CAAACCATCAGCGTTCGCGCTTATTGGAGTTTT |
| L2288A-For | ATCAACGCTGATGGTGCGTACGAGGTTATACTT |
| L2288A-Rev | AAGTATAACCTCGTACGCACCATCAGCGTTGAT |
| V2291A-For | GATGGTTTGTACGAGGCGATACTTCAAGGAGTA |
| V2291A-Rev | TACTCCTTGAAGTATCGCCTCGTACAAACCATC |
| Y2299A-For | CAAGGAGTATATACTGCGCCATACCATGGCGAT |
| Y2299A-Rev | ATCGCCATGGTATGGCGCAGTATATACTCCTTG |
| H2302A-For | TATACTTATCCATACGCGGGCGATGGTGTTTGT |
| H2302A-Rev | ACAAACACCATCGCCCGCGTATGGATAAGTATA |
| D2304A-For | TATCCATACCATGGCGCGGGTGTTTGTGGTTCG |
| D2304A-Rev | CGAACCACAAACACCCGCGCCATGGTATGGATA |
| C2307A-For | CACGGCGATGGTGTTGCGGGTTCGATATTGTTG |

| | |
|---|---|
| C2307A-Rev | CAACAATATCGAACCCGCAACACCATCGCCGTG |
| H2324A-For | TCAGTACCAGCAACAGCGATACCTATAATTGG |
| H2324A-Rev | CCAATTATAGGTATCGCTGTTGCTGGTACTGA |
| E2329A-For | CATGTTGCTGGTACTGCGGGATTGCATGGCTTT |
| E2329A-Rev | AAAGCCATGCAATCCCGCAGTACCAGCAACATG |
| Q2118A-For | GTGACTACTAAGCCTGCGGGATCAACACAACAA |
| Q2118A-Rev | TTGTTGTGTTGATCCCGCAGGCTTAGTAGTCAC |
| E2180A-For | ACTGCCGCCTTTCCTGCGGGAACCAAGTACTAT |
| E2180A-Rev | ATAGTACTTGGTTCCCGCAGGAAAGGCGGCAGT |

A

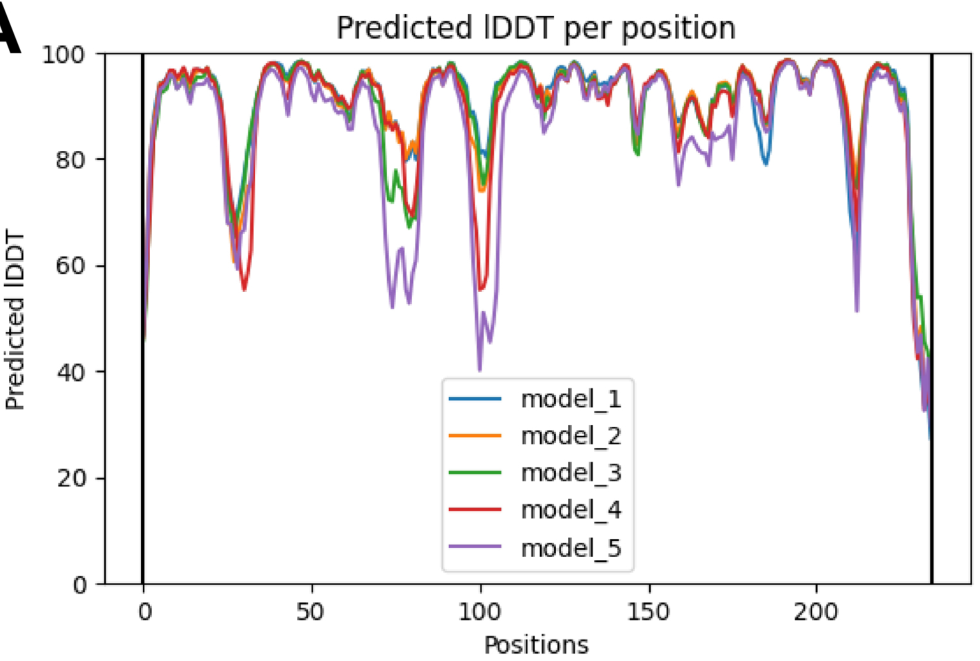

Predicted lDDT per position
Predicted lDDT
Positions
100
80
60
40
20
0
0
50
100
150
200
model_1
model_2
model_3
model_4
model_5


B

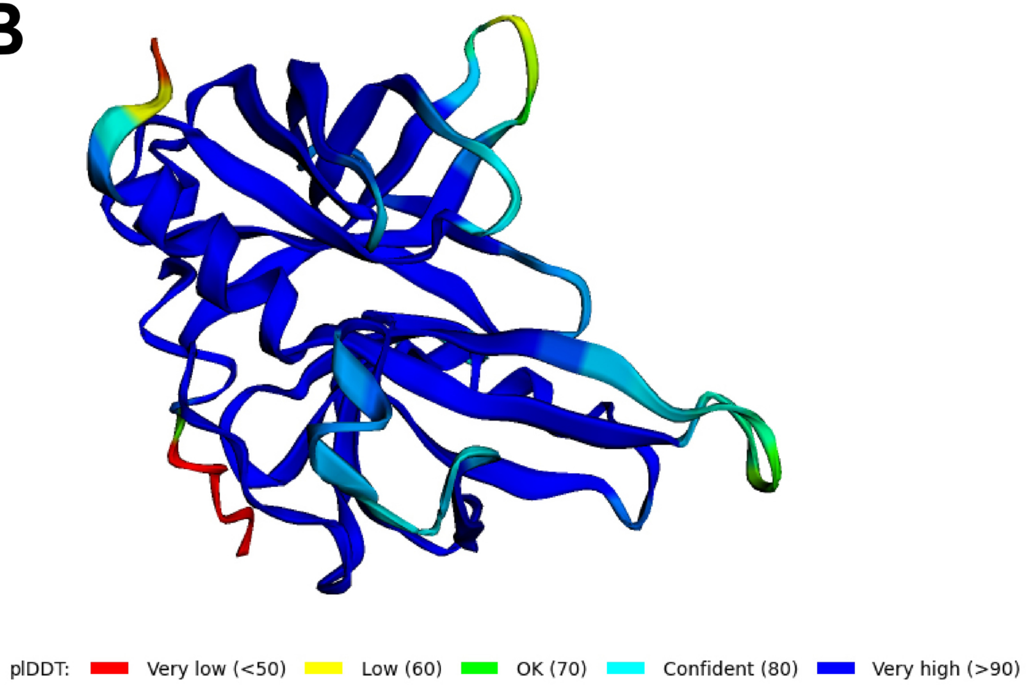

plDDT:
Very low (<50)
Low (60)
OK (70)
Confident (80)
Very high (>90)

```
Bemisia             --------------------------------DAVQRKVVANTVIFTITS---DQTD-RT
Brevicoryne         ------------PQSSV-------SNNNLELISSVYNRINRNTVYI--NYED-KVLN-KI
Aedes               -------------------------EGPKSSLDGVLNKISSNTVIVIGKYTE-DGVE-RI
Ista                IKASQSIRAKALRLVKPEMAQRESLQEVQNQFEEKLKIIQGNTIFLIGVDPI-SGV---T
Culex               -----------------------------------VDKILGNTVFIIARHPE-TGA---V
Hypera              --------------------------NATNQEEVISAKIIRNTFFLYAEYGD-ETRM-TS
Watson              -----TIGAKANKKIPIAKPK---AQACAQQEDVIFKYIKRNTFWLSLEFET-ETGY-DS
Serbia              ------------------VPVKALAQSCEQQENVITKLIMKNTFWILAEYVE-HGII-KT
Rondonia            -------ACKVAKHPIARPEIAPKFQSTSQQSRELPRFLEENMVSIHADFEDERGIT-RT
Antheraea           -SKAKVTKAPKAKVRTPVRATSKLEYQSAQQFEVVQQRLKDNMSAIDVVYTNADGKD-VR
Lymantria           ------------------RATKTLEYQGAQQFDVVKQRLRNNLSSIDVVYTDVEGNM-KR
Moku                ----YCKIDKAKPAKANFKPADKEDDGSLDRIEQLERKLLNNICFLEARWNDEKGDA-KM
Moran               -------KSPRAGRKPDAPA----QKESAENDYYVERKLLNNVCFLEFDYINSDNKR-RC
Pityohyphantes      ---------------------------ANQNVGDVMRKISNNIVFLQLSYTW-EGIP-KS
Tribolium           ------------VKRPDVKVMSKMPEMAPQQYSNIMAAINRNTVIITAEYFNEKGSL-CS
Nilaparvata         ---------------------------SSQQSEAVIKIIRNNTFYLSICD---NTTM-RI
Erysiphe            -------------VARTAPRIGAKAQGNSQQAEYVEGLIRKNMVILRVRGTG-ELVG-RV
Psammotettix        -----TPKHPVKSKVPISSKTFVTPQAGAIQIAQLSNKITFNTVFVVVQYEC-EGRL-LE
Euscelidius         -------KFPQKPKVPTSEKVFNPPEMSTEQRQVMINRINKNGVLIYVNWVQ-DGTR-MN
Formica             -NLRYDYQTPIKNPPKHIPVAFAKPQSNLPQIEVVTKLIHRNTFFLHCYIDD----R-HI
King                --------APKAGVKPVVS------QSESQQYEVVLKLIRSNFFYLTMV-----ANG-KV
Pittosporum         -------------------------QAMMPQYNACIDRINKNSCTVVCYYGV--NSD-KM
Darwin              -GLKYGEHVTPKARLSPRNVPSAKPQDDTQQVDVALKRIRANMCYIECHRPD----G-YT
DWV                 -GLKYSEAVTVKAPRIHRLPVTTKPQGSTQQVDAAVNKILQNMVYIGVVFPKVPGSKWRD
                                                          :  *   .

Bemisia             ARCFGLGIYNRFVLLPKHYKLTFLNL-----TTEKLYCEPFGERHLRQHIDPT-------
Brevicoryne         SHMKCIVIKERYVIVLRHYIEYLECNAT---DTSNVNITWADCGSY-----------PFN
Aedes               ARARCLALHTRTLLVLRHYFEFFKYK-----NVEKVTVVSRKGDCSV-EYGLDEL--QFM
Ista                YKGRCLGLYDRTVLAIKHYFDHFLHK-----GIKRLKVVRPNSNTTIYDVDIDNL--KVQ
Culex               YKGRCLGLYGKEVLVVKHYFDLFRKI-----GVKKVEIVRPRVEATIFTVDISDL--LFA
Hypera              RVFRCLGLNRHNFIIIDHYVTFLRQQ-----PNLKLSFVRDGCCVEISTA----------
Watson              RLFRCLGLVGKYFLFLDHYYNFMKSKP----NAKLFYIQQGMYIEIPLSVVNQNI-----
Serbia              KILRCLGLYKHYFLLLDHYIGALRMMD----DLKLSFIKQGTYITIPNSVVFTNI-----
Rondonia            SVSSCFVICDRRMLILRHYWDHWIKLP----PTTKFYLVSRKTVSVNY---PKGI--PLT
Antheraea           VRNYGLMLRDQQMLIQKHYYDFWRRLD----VTAKFYF----CNQKFKSIHPDGI--IIS
Lymantria           TRNFGLMLKDQQMLIQKHYYDFWKRLD----LTAKFYFYNNNIKSHA----PDGI--LLT
Moku                LYGRCLGIRERQVLILKHYIEEMLHIP---SDATFMINYIMSGKPCT----------GFL
Moran               ITGRCLGIRNREIVVIKHYLEEMKSLAKKYPEHGLTIVLCRNELDSRIKLTMEDLD-KVS
Pityohyphantes      KKFRVVMLRERQCLMIRHYIEDIQFYADHD-PNAKLSLLVNGNNEVPLAINCVN---EFS
Tribolium           MKARALGLEANRVIMIRHYHDEFTAMP----PSTRYYVSLVTNNQLRPRV-------EIN
Nilaparvata         YRYRCLGIHGHFALMLRHYVDNIKDKIRCV-GTTNLTIAVEYNHNCVLTG-SRTI--QLD
Erysiphe            WNFVCICLGDRNILALRHYLDVIDRYS----NSVTLSLAYAERHDSRLKV-LDGV--EML
Psammotettix        KSCRCLMLRGRAMLILRHYWEEYQYLVD---RGYKLDVALHFGAGRK-SAEPRHVIRRVG
Euscelidius         RNCRCLMLGGRNMLVLRHYLEEYSALVE---QGFKLNVDLVF--GNK-IGDPVKV--NIT
Formica             VAMRGVVMANREALVLRHYIEQIKGLQERYKEHLKITFTWHQQQYERMNC--KEL--EID
King                VTYRCLRLRMNEVLILRHYDDEIR-------SVGDVAVTYGKCAGLNADTFKYGI--QFN
Pittosporum         IHFRCLILRNREMLFLRHYREQVVNL-----PNAQFRFKFNVDNEKR-TGNPDGI--PFD
Darwin              KRYRALLLKNRMMLFLRHYHENIMKQP----EGTLVIFTYAFNSEERIKC-VRGI--ELR
DWV                 INFRCLMLHNRQCLMLRHYIESTAAFP----EGTKYYFKYIHNQETRMSGDISGI--EID
                         . :  .  :   **
```

```
Bemisia             -WL-VY--------------------EDIEDADLCILRLPANFP-MFKDIRKYMATMHDH
Brevicoryne         -YK-DY--KIYWCE---------------NSN-IGVLRLGNWFS-ARRSLIPFITKSDSF
Aedes               -WSDDC--------------------G------YGTCELPQSYPVNFKKITQLICSDKFS
Ista                -WG-DC--------------------G------YGLLYLPQSYPTQFRKITQFFPSEEQT
Culex               -WG-DC--------------------G------YGMMTLPRSFPTQFKKITQLIASEELN
Hypera              -FLSEC--------------------KAIKDSALMVAEMDKTVP-QFVNIIKFFIRSNQT
Watson              --------------------------QTLTNSTFMIGKMHKTIP-HFANIIKLMIKQEAS
Serbia              --------------------------KKLQNSALCIGTMYKTIP-MFSNIIKFIIKAGDT
Rondonia            -LG-NL--QPEWFADIDSS-------GFYKSH-FGIAQLPQSSP-AFKDLRKFIATVEDH
Antheraea           NLF-DL--DVDWFMTPGL--------EYYDSN-FGVLHLPKTVP-AFKDITRFIAKSSEH
Lymantria           NFF-DL--DVDWFMTPDQ--------DIFDSN-FGILHLPKIVP-AYKDLTKFIAKSTEH
Moku                -TR-KCLESVTYFSIN----------GKINASNYGILTLPKFMP-MFKDILNSIVRKNDH
Moran               -WM-KIG-------------------NEENTSNFGILTLPIRVP-QFRNIVNSIATIGQH
Pityohyphantes      -VL-ENQIETDFDEEREGT-------GYVLYNGLCVVDLPKTVR-EFPTITRHFVSAKDE
Tribolium           -YR-ECRCDAFYCKD-----------PKYGTN-FFMLYMPPHIP-LFRNIKNLIPTAESH
Nilaparvata         -ND-AFLASCKFFKHKAPEFYSEDQFNAHQSN-FVVFKVPSQCL-SFKSLLKFFPSQAEV
Erysiphe            -YE-NLHRQEYFYDE-----------GKIDSN-FVVFQAPARFQ-PMRNLMKFIASGQEH
Psammotettix        -WA-EL-QKVAWCSS--SS-------GELTSN-FGICELPMYIP-MFCDITNYIASMSEH
Euscelidius         -FT-ELMSQVAYCDN---------------SN-FCIVVLPKFVR-QFPKIYPYFATRANH
Formica             -FL-NC--RIIYYGSTDLS-------GCAMSN-LCIVELPARIP-EXXXLKKFVCAEASH
King                -YR-EC--KIKWYRNVVVD-------GCYPSN-FGILLLPPCFP-MAKDLTKFIATSDDH
Pittosporum         -FL-GC--KTSYYGSVE---------NCMFTSNLGIMVLPVNIP-ECRNITKFIMPLNKH
Darwin              -YQ-DL--EKTEYG---RQ-------DQYDSN-FMIVKLPASIP-EAKDLTKFISPIDEH
DWV                 -LL-NL--PRLYYGGLAGE-------ESFDSN-IVLVTMPNRIP-ECKSIIKFIASHNEH
                                                                     :   .

Bemisia             EVRGLPRQAYILSPPIRKAPHDIFKEIAVNISGYY------D-DLSTKCDGVQIVT---K
Brevicoryne         LGNAGRECVILDVK------LSESYVYNENIKLVD------N-VTIAPTSYSKMLIM--P
Aedes               RCYP-SNMMIVEHN------FSNLHTLEVQAKVMS------ESKKIPQQPGMSSWVI--H
Ista                SEYP-ASVQMFEDS------GAGYKLFSMEMTVID------E-VVVPKSGTFEAWTI--S
Culex               TEYP-AAIQIVEDK------GVGHTLISTNMRLIQ------E-QVVPANQEFEAWTI--F
Hypera              GNLS-PLATLHEYVWDANKQNYSLRSRDFSRISRQ------N-CVSVTNDDDSRTIV--S
Watson              GNIA-PQAKLYEYK-LNSDREFMMQIHHIDRIYRQ------D-RLEVNNDDGSVSRI--G
Serbia              KNIC-PQAKLYEYS--VEGESYVMKVQSFARIMRK------D-QVDVSNDDNTVTHV--S
Rondonia            RYIDTQACYLYDSQ--------KRCQVILPIQLQK------G-YIVSDGDNGTHLPI---
Antheraea           SYIKFDECYLYSSL------TDMCMHCVMNVEMNK------D---VTDSNGWLQL----S
Lymantria           QYIKFDECYLYSSL------SGESMHCVMNIEYNK------E---VTDANGWLRL----D
Moku                HYVA-SQGHIMSSD----MEGRILRHRHMQLRPRA------F-LVIDGDDEGHVSTICND
Moran               LNVR-SVGDFLSHK------GKTRRRIVIQEKRV---------LVVSKTENTTGAIL--D
Pityohyphantes      LRMS-NNGILLMPEPGKGSLANTIVCENVKFSNYE------N-LTVNAADTTSAVHL--E
Tribolium           QYCG-RLGHLYAFG------DATYADLAFNFEG--------E-YRIQQWGNAPGMVC--M
Nilaparvata         NQAS-TTLRLVNEK------DN--MELYGKLKSHKVGGLYIP-PVHKENAYNDAVTM--A
Erysiphe            TYIS-QKCMFIRPN------QLATEVCVETRNMPV---------IIEPACEGAAEVHL--V
Psammotettix        ENVS-SLCDLYVVN------GE--SKFGMPLSVKR------R-FAVAETASSSQVKV--D
Euscelidius         DNVS-GKVDMITVA------GE--SSFDLPVSVSK------N-FVVSETANSSEVVC--E
Formica             CRAG-TTGRLIEPG------THNILQLKIDYNSKM------P-YVIEPYGDTKQVNM--L
King                KYSS-GFGSLVEGD-------KVVEHMRITRDSKR------P-FVIPKTDIVSPVAM--D
Pittosporum         RYAS-TTGSLVQPG------GNTLNQVTIDYSNKQ------P-HTIAGMNGSTDVYM--Q
Darwin              RRTL-SDGILVNEK------NTELVHFS---PQNR------P-HIIAEGD-LRDVIM--E
DWV                 IRAQ-NDGVLVTGD------HTQLLAFE---NNNK------T-PISINADGLYEVIL--Q
```

```
Bemisia             EAIQYNY-SQPGMCGAILLSRNTQR---PILGMHVAGTCVDF-GFQGMGFSAILVQEIF-
Brevicoryne         NAYMYKK-NYPGMCGSVLMNEATNT---PILGIHVAGAN----GK---GFSEPICRAQF-
Aedes               DGFEYAW-GGKGKCGSLLLVPSLSS---PIVGIHSAGVKD---VR---GFAEVLFRETF-
Ista                NGFKYPW-GGSGRCGSLLLSPKLAS---PIIGIHTAGTD----GRI--GYAERLLRETF-
Culex               DGFKYDW-GGSGCCGSLILSPTLSS---PIIGIHTAGIGK---TT---GFGERLYRESFD
Hypera              CLYEYPY-GGKGVCGSVLVSRQPLT--NPILGIHIAGFSD---GSR--GYSEAIVYETF-
Watson              ANYEYPV-SGRGVCGSVLVSSSNMN--APILGIHVAGIKN---GIE--GYAEALVYESFS
Serbia              SLYAYPY-AHRGVCGSVLVSTSNMN--APILGIHIAGFSN---GVE--GFAEAITYETF-
Rondonia            -AYKYRY-TGDGKCGSILVCGNLQR---PIIGMHFAGSN----IW---GSSEPIYSENFK
Antheraea           ECYSYKY-SAKGLCGSALLCATLER---PIIGIHFAGTK----VF---GYAEPIAYETF-
Lymantria           ECYSYKY-TGVGLCGSALLCSTLER---PIIGVHFAGTS----TY---GYAEPLCYESF-
Moku                GAYEYSV-HGKGMCGSLLISDKVCRGNPGIIGLHVAGSK----GN---GYAEPIYREMF-
Moran               MAYQYGH-HGQGLCGSLLVCENVCNGNIGIIGMHVAGRS----GI---GYSEPLCREWFT
Pityohyphantes      RVWMYRGVHGYGLCGSLLLNEDSGK----IIGIHTAGSSTN--NV---GFSERLVYEEW-
Tribolium           NAYTYSK-HKPGLCGSVLISEGLPS--TPLIGMHVAGAN----GK---GYAEPLFREMF-
Nilaparvata         SYWEYGV-HGRGMCGSVLVSNNLQN---PIVGMHVAGCDNA--IK---GYSELIVKEMF-
Erysiphe            EVYKYPKMQREGFCGSIIICPTLQC---PIIGMHVAGHSEGM-GY---GIAEPLVKEMF-
Psammotettix        RAYQYNH-QYKGLCGSVLVSRKGNSGNGSIIGIHVAGSQSS--GN---GFSEPIVREYLD
Euscelidius         RVYGYAK-RSPGLCGSVIISNSLGSGNGAIIGMHIAGNAAS--GT---GYAEPLYQEMFA
Formica             NVYTYDY-HAQGACGSILLADNLEQ---PIVGIHVAGTVGKGCGR---GVAEPLSAEMM-
King                DVYVYTK-HGAGVCGSVLLAANMQR---PIIGMHVAGYGTVS-GN---GLAEILCSEMFS
Pittosporum         TVYTYQY-HAPGMCGSVLLSSGSET---PIIGIHVAGYEGSGMSY---GMSEILHREMFD
Darwin              TSYTYNY-HGRGVCGSVLLSRNLQK---PIIGMHVAGVEGVN-GY---GISEPLFYETFV
DWV                 GVYTYPY-HGDGVCGSILLSRNLQR---PIIGIHVAGTEGLH-GF---GVAEPLVHEMFT
                        *      * **: ::          ::*:* **           * .  :

Bemisia             --------
Brevicoryne         --------
Aedes               --------
Ista                --------
Culex               ADSESE--
Hypera              --------
Watson              --------
Serbia              --------
Rondonia            RSVEKE--
Antheraea           --------
Lymantria           --------
Moku                --------
Moran               G-------
Pityohyphantes      --------
Tribolium           --------
Nilaparvata         --------
Erysiphe            --------
Psammotettix        G-------
Euscelidius         --------
Formica             --------
King                --------
Pittosporum         NMPLPEE-
Darwin              SVPKSIDT
DWV                 GKAIESE-
```

>DWV
GLKYSEAVTVKAPRIHRLPVTTKPQGSTQQVDAAVNKILQNMVYIGVVFPKVPGSKWRDINFRCLMLHNRQCLMLRHYIE
STAAFPEGTKYYFKYIHNQETRMSGDISGIEIDLLNLPRLYYGGLAGEESFDSNIVLVTMPNRIPECKSIIKFIASHNEH
IRAQNDGVLVTGDHTQLLAFENNNKTPISINADGLYEVILQGVYTYPYHGDGVCGSILLSRNLQRPIIGIHVAGTEGLHG
FGVAEPLVHEMFTGKAIESE

>QCL11100.1:2261-2493 polyprotein [King virus]
APKAGVKPVVSQSESQQYEVVLKLIRSNFFYLTMVANGKVVTYRCLRLRMNEVLILRHYDDEIRSVGDVAVTYGKCAGLN
ADTFKYGIQFNYRECKIKWYRNVVVDGCYPSNFGILLLPPCFPMAKDLTKFIATSDDHKYSSGFGSLVEGDKVVEHMRIT
RDSKRPFVIPKTDIVSPVAMDDVYVYTKHGAGVCGSVLLAANMQRPIIGMHVAGYGTVSGNGLAEILCSEMFS

>AWK77848.1:2012-2262 polyprotein [Darwin bee virus 3]
GLKYGEHVTPKARLSPRNVPSAKPQDDTQQVDVALKRIRANMCYIECHRPDGYTKRYRALLLKNRMMLFLRHYHENIMKQ
PEGTLVIFTYAFNSEERIKCVRGIELRYQDLEKTEYGRQDQYDSNFMIVKLPASIPEAKDLTKFISPIDEHRRTLSDGIL
VNEKNTELVHFSPQNRPHIIAEGDLRDVIMETSYTYNYHGRGVCGSVLLSRNLQKPIIGMHVAGVEGVNGYGISEPLFYE
TFVSVPKSIDT

>QED21536.1:2477-2716 polyprotein [Moran virus]
KSPRAGRKPDAPAQKESAENDYYVERKLLNNVCFLEFDYINSDNKRRCITGRCLGIRNREIVVIKHYLEEMKSLAKKYPE
HGLTIVLCRNELDSRIKLTMEDLDKVSWMKIGNEENTSNFGILTLPIRVPQFRNIVNSIATIGQHLNVRSVGDFLSHKGK
TRRRIVIQEKRVLVVSKTENTTGAILDMAYQYGHHGQGLCGSLLVCENVCNGNIGIIGMHVAGRSGIGYSEPLCREWFTG

>YP_009351892.1:2274-2510 polyprotein [Pityohyphantes rubrofasciatus iflavirus]
ANQNVGDVMRKISNNIVFLQLSYTWEGIPKSKKFRVVMLRERQCLMIRHYIEDIQFYADHDPNAKLSLLVNGNNEVPLAI
NCVNEFSVLENQIETDFDEEREGTGYVLYNGLCVVDLPKTVREFPTITRHFVSAKDELRMSNNGILLMPEPGKGSLANTI
VCENVKFSNYENLTVNAADTTSAVHLERVWMYRGVHGYGLCGSLLLNEDSGKIIGIHTAGSSTNNVGFSERLVYEEW

>QNS17457.1:2158-2380 RNA-dependent RNA polymerase [Serbia picorna-like virus 2]
VPVKALAQSCEQQENVITKLIMKNTFWILAEYVEHGIIKTKILRCLGLYKHYFLLLDHYIGALRMMDDLKLSFIKQGTYI
TIPNSVVFTNIKKLQNSALCIGTMYKTIPMFSNIIKFIIKAGDTKNICPQAKLYEYSVEGESYVMKVQSFARIMRKDQVD
VSNDDNTVTHVSSLYAYPYAHRGVCGSVLVSTSNMNAPILGIHIAGFSNGVEGFAEAITYETF

>QKK82957.1:1131-1364 hypothetical protein [Pittosporum tobira picorna-like virus]
QAMMPQYNACIDRINKNSCTVVCYYGVNSDKMIHFRCLILRNREMLFLRHYREQVVNLPNAQFRFKFNVDNEKRTGNPDG
IPFDFLGCKTSYYGSVENCMFTSNLGIMVLPVNIPECRNITKFIMPLNKHRYASTTGSLVQPGGNTLNQVTIDYSNKQPH
TIAGMNGSTDVYMQTVYTYQYHAPGMCGSVLLSSGSETPIIGIHVAGYEGSGMSYGMSEILHREMFDNMPLPEE

>QHD64830.1:2097-2334 RdRp [Erysiphe necator associated picorna-like virus 1]
VARTAPRIGAKAQGNSQQAEYVEGLIRKNMVILRVRGTGELVGRVWNFVCICLGDRNILALRHYLDVIDRYSNSVTLSLA
YAERHDSRLKVLDGVEMLYENLHRQEYFYDEGKIDSNFVVFQAPARFQPMRNLMKFIASGQEHTYISQKCMFIRPNQLAT
EVCVETRNMPVIIEPACEGAAEVHLVEVYKYPKMQREGFCGSIIICPTLQCPIIGMHVAGHSEGMGYGIAEPLVKEMF

>QQL13637.1:2182-2425 polyprotein [Antheraea mylitta iflavirus]
SKAKVTKAPKAKVRTPVRATSKLEYQSAQQFEVVQQRLKDNMSAIDVVYTNADGKDVRVRNYGLMLRDQQMLIQKHYYDF
WRRLDVTAKFYFCNQKFKSIHPDGIIISNLFDLDVDWFMTPGLEYYDSNFGVLHLPKTVPAFKDITRFIAKSSEHSYIKF
DECYLYSSLTDMCMHCVMNVEMNKDVTDSNGWLQLSECYSYKYSAKGLCGSALLCATLERPIIGIHFAGTKVFGYAEPIA
YETF

>YP_009047245.1:2212-2438 polyprotein [Lymantria dispar iflavirus 1]
RATKTLEYQGAQQFDVVKQRLRNNLSSIDVVYTDVEGNMKRTRNFGLMLKDQQMLIQKHYYDFWKRLDLTAKFYFYNNNI
KSHAPDGILLTNFFDLDVDWFMTPDQDIFDSNFGILHLPKIVPAYKDLTKFIAKSTEHQYIKFDECYLYSSLSGESMHCV
MNIEYNKEVTDANGWLRLDECYSYKYTGVGLCGSALLCSTLERPIIGVHFAGTSTYGYAEPLCYESF

>QHI42120.1:2186-2431 polyprotein [Rondonia iflavirus 1]
ACKVAKHPIARPEIAPKFQSTSQQSRELPRFLEENMVSIHADFEDERGITRTSVSSCFVICDRRMLILRHYWDHWIKLPP
TTKFYLVSRKTVSVNYPKGIPLTLGNLQPEWFADIDSSGFYKSHFGIAQLPQSSPAFKDLRKFIATVEDHRYIDTQACYL
YDSQKRCQVILPIQLQKGYIVSDGDNGTHLPIAYKYRYTGDGKCGSILVCGNLQRPIIGMHFAGSNIWGSSEPIYSENFK
RSVEKE

>YP_009305421.1:2255-2502 polyprotein [Moku virus]
YCKIDKAKPAKANFKPADKEDDGSLDRIEQLERKLLNNICFLEARWNDEKGDAKMLYGRCLGIRERQVLILKHYIEEMLH
IPSDATFMINYIMSGKPCTGFLTRKCLESVTYFSINGKINASNYGILTLPKFMPMFKDILNSIVRKNDHHYVASQGHIMS
SDMEGRILRHRHMQLRPRAFLVIDGDDEGHVSTICNDGAYEYSVHGKGMCGSLLISDKVCRGNPGIIGLHVAGSKGNGYA
EPIYREMF

>YP_009328891.1:2309-2548 polyprotein [Euscelidius variegatus virus 1]
KFPQKPKVPTSEKVFNPPEMSTEQRQVMINRINKNGVLIYVNWVQDGTRMNRNCRCLMLGGRNMLVLRHYLEEYSALVEQ
GFKLNVDLVFGNKIGDPVKVNITFTELMSQVAYCDNSNFCIVVLPKFVRQFPKIYPYFATRANHDNVSGKVDMITVAGES
SFDLPVSVSKNFVVSETANSSEVVCERVYGYAKRSPGLCGSVIISNSLGSGNGAIIGMHIAGNAASGTGYAEPLYQEMFA

>AUE23905.1:2261-2492 polyprotein, partial [Tribolium castaneum iflavirus]
VKRPDVKVMSKMPEMAPQQYSNIMAAINRNTVIITAEYFNEKGSLCSMKARALGLEANRVIMIRHYHDEFTAMPPSTRYY
VSLVTNNQLRPRVEINYRECRCDAFYCKDPKYGTNFFMLYMPPHIPLFRNIKNLIPTAESHQYCGRLGHLYAFGDATYAD
LAFNFEGEYRIQQWGNAPGMVCMNAYTYSKHKPGLCGSVLISEGLPSTPLIGMHVAGANGKGYAEPLFREMF

>YP_008130310.1:2380-2618 polyprotein [Nilaparvata lugens honeydew virus-3]
SSQQSEAVIKIIRNNTFYLSICDNTTMRIYRYRCLGIHGHFALMLRHYVDNIKDKIRCVGTTNLTIAVEYNHNCVLTGSR
TIQLDNDAFLASCKFFKHKAPEFYSEDQFNAHQSNFVVFKVPSQCLSFKSLLKFFPSQAEVNQASTTLRLVNEKDNMELY
GKLKSHKVGGLYIPPVHKENAYNDAVTMASYWEYGVHGRGMCGSVLVSNNLQNPIVGMHVAGCDNAIKGYSELIVKEMF

>YP_009553259.1:2363-2614 polyprotein [Psammotettix alienus iflavirus 1]
TPKHPVKSKVPISSKTFVTPQAGAIQIAQLSNKITFNTVFVVVQYECEGRLLEKSCRCLMLRGRAMLILRHYWEEYQYLV
DRGYKLDVALHFGAGRKSAEPRHVIRRVGWAELQKVAWCSSSSGELTSNFGICELPMYIPMFCDITNYIASMSEHENVSS
LCDLYVVNGESKFGMPLSVKRRFAVAETASSSQVKVDRAYQYNHQYKGLCGSVLVSRKGNSGNGSIIGIHVAGSQSSGNG
FSEPIVREYLDG

>QGA70909.1:2053-2286 RNA-dependent RNA polymerase [Ista virus]
IKASQSIRAKALRLVKPEMAQRESLQEVQNQFEEKLKIIQGNTIFLIGVDPISGVTYKGRCLGLYDRTVLAIKHYFDHFL
HKGIKRLKVVRPNSNTTIYDVDIDNLKVQWGDCGYGLLYLPQSYPTQFRKITQFFPSEEQTSEYPASVQMFEDSGAGYKL
FSMEMTVIDEVVVPKSGTFEAWTISNGFKYPWGGSGRCGSLLLSPKLASPIIGIHTAGTDGRIGYAERLLRETF

>QWC36469.1:1934-2146 polyprotein [Bemisia tabaci ifla-like virus 1]
DAVQRKVVANTVIFTITSDQTDRTARCFGLGIYNRFVLLPKHYKLTFLNLTTEKLYCEPFGERHLRQHIDPTWLVYEDIE
DADLCILRLPANFPMFKDIRKYMATMHDHEVRGLPRQAYILSPPIRKAPHDIFKEIAVNISGYYDDLSTKCDGVQIVTKE
AIQYNYSQPGMCGAILLSRNTQRPILGMHVAGTCVDFGFQGMGFSAILVQEIF

>QED21508.1:2165-2399 polyprotein [Watson virus]
TIGAKANKKIPIAKPKAQACAQQEDVIFKYIKRNTFWLSLEFETETGYDSRLFRCLGLVGKYFLFLDHYYNFMKSKPNAK
LFYIQQGMYIEIPLSVVNQNIQTLTNSTFMIGKMHKTIPHFANIIKLMIKQEASGNIAPQAKLYEYKLNSDREFMMQIHH
IDRIYRQDRLEVNNDDGSVSRIGANYEYPVSGRGVCGSVLVSSSNMNAPILGIHVAGIKNGIEGYAEALVYESFS

>QUS52852.1:2169-2384 polyprotein [Hypera postica associated iflavirus 1]
NATNQEEVISAKIIRNTFFLYAEYGDETRMTSRVFRCLGLNRHNFIIIDHYVTFLRQQPNLKLSFVRDGCCVEISTAFLS
ECKAIKDSALMVAEMDKTVPQFVNIIKFFIRSNQTGNLSPLATLHEYVWDANKQNYSLRSRDFSRISRQNCVSVTNDDDS
RTIVSCLYEYPYGGKGVCGSVLVSRQPLTNPILGIHIAGFSDGSRGYSEAIVYETF

>QRW42579.1:2196-2401 polyprotein [Culex Iflavi-like virus 3]
VDKILGNTVFIIARHPETGAVYKGRCLGLYGKEVLVVKHYFDLFRKIGVKKVEIVRPRVEATIFTVDISDLLFAWGDCGY
GMMTLPRSFPTQFKKITQLIASEELNTEYPAAIQIVEDKGVGHTLISTNMRLIQEQVVPANQEFEAWTIFDGFKYDWGGS
GCCGSLILSPTLSSPIIGIHTAGIGKTTGFGERLYRESFDADSESE

>YP_001285409.1:2210-2424 polyprotein [Brevicoryne brassicae virus - UK]
PQSSVSNNNLELISSVYNRINRNTVYINYEDKVLNKISHMKCIVIKERYVIVLRHYIEYLECNATDTSNVNITWADCGSY
PFNYKDYKIYWCENSNIGVLRLGNWFSARRSLIPFITKSDSFLGNAGRECVILDVKLSESYVVYNENIKLVDNVTIAPTSY
SKMLIMPNAYMYKKNYPGMCGSVLMNEATNTPILGIHVAGANGKGFSEPICRAQF

>QGW51140.1:2015-2226 polyprotein [Aedes vexans iflavirus]
EGPKSSLDGVLNKISSNTVIVIGKYTEDGVERIARARCLALHTRTLLVLRHYFEFFKYKNVEKVTVVSRKGDCSVEYGLD
ELQFMWSDDCGYGTCELPQSYPVNFKKITQLICSDKFSRCYPSNMMIVEHNFSNLHTLEVQAKVMSESKKIPQQPGMSSW
VIHDGFEYAWGGKGKCGSLLLVPSLSSPIVGIHSAGVKDVRGFAEVLFRETF

>AWI42879.1:2083-2335 polyprotein, partial [Formica cinerea virus 2]
NLRYDYQTPIKNPPKHIPVAFAKPQSNLPQIEVVTKLIHRNTFFLHCYIDDRHIVAMRGVVMANREALVLRHYIEQIKGL
QERYKEHLKITFTWHQQQYERMNCKELEIDFLNCRIIYYGSTDLSGCAMSNLCIVELPARIPEXXXLKKFVCAEASHCRA
GTTGRLIEPGTHNILQLKIDYNSKMPYVIEPYGDTKQVNMLNVYTYDYHAQGACGSILLADNLEQPIVGIHVAGTVGKGC
GRGVAEPLSAEMM

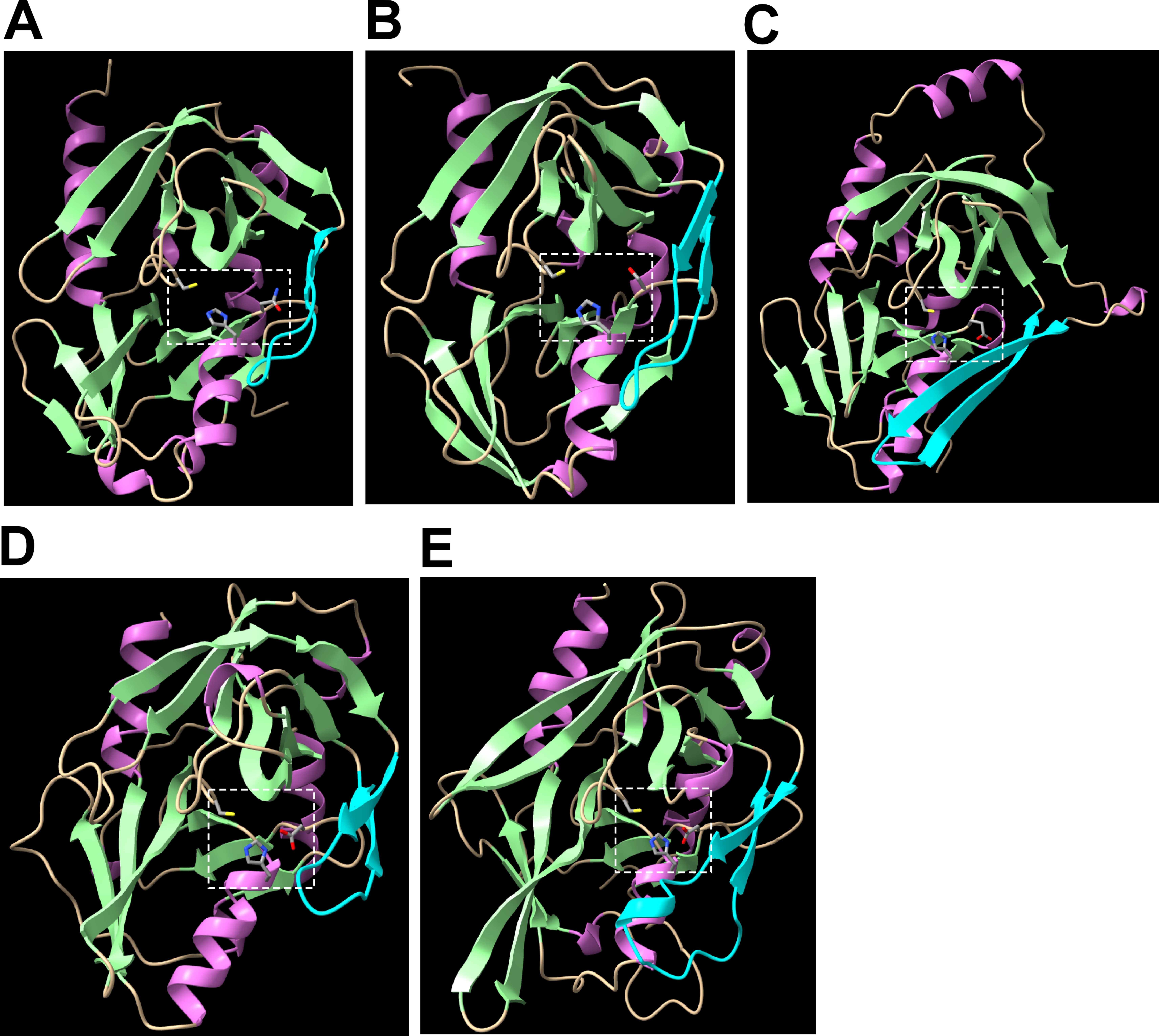
A
B
C
D
E

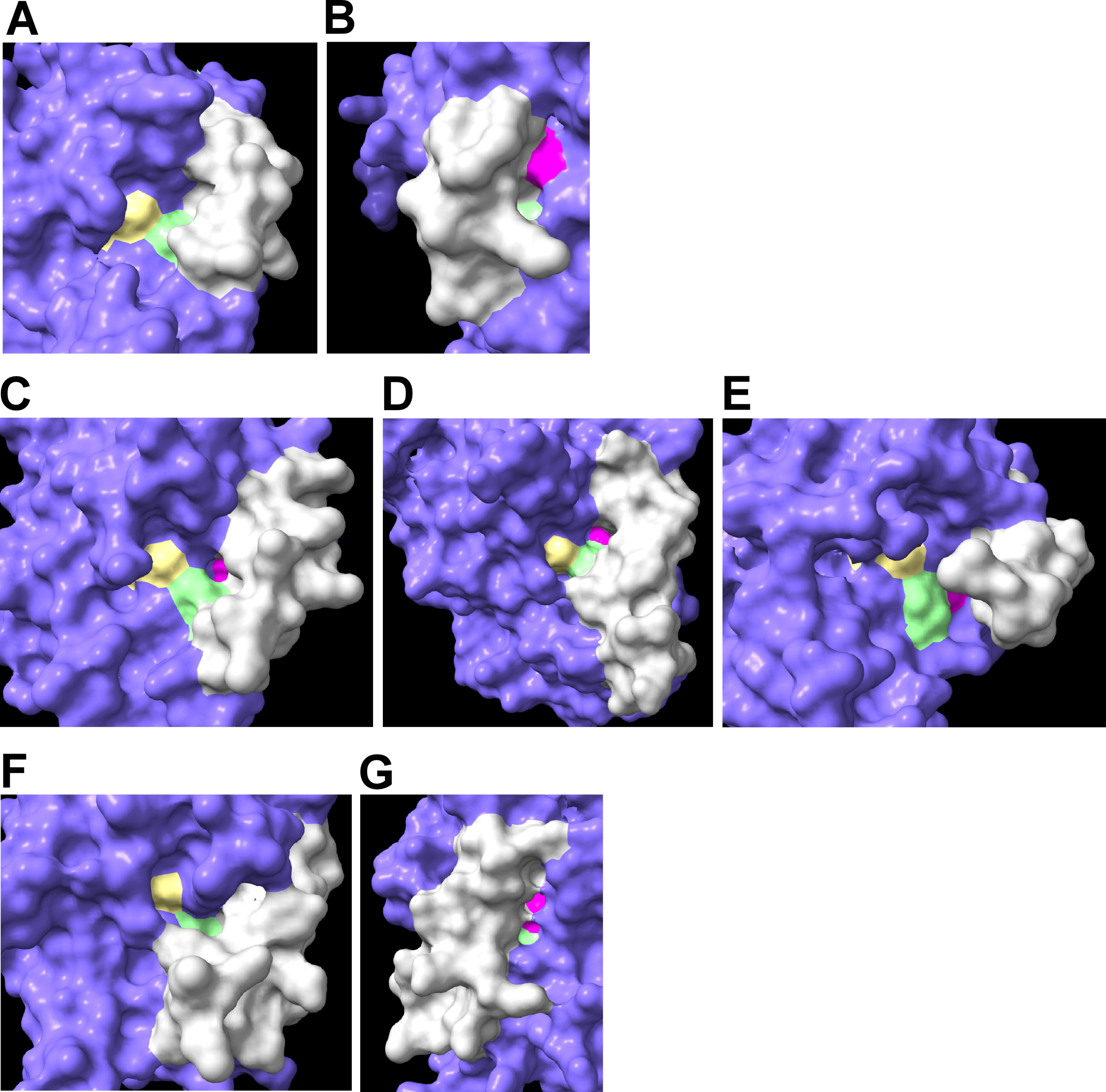
